# Volumetric refractive index measurement and quantitative density analysis of mouse brain tissue with sub-micrometer spatial resolution

**DOI:** 10.1101/2023.01.16.524195

**Authors:** Ariel J. Lee, Young Seo Kim, Joon-Goon Kim, Herve Hugonnet, Moosung Lee, Taeyun Ku, YongKeun Park

**Affiliations:** Department of Physics, Korea Advanced Institute of Science and Technology (KAIST), Daejeon, 34141, Republic of Korea; Graduate School of Medical Science and Engineering, KAIST, Daejeon, 34141, Republic of Korea; Tomocube Inc., Daejeon, 34109, Republic of Korea

**Keywords:** microscopy, tissue, three-dimensional imaging, refractive index, quantitative phase imaging

## Abstract

High-resolution structural imaging of brain tissue is important for neuroscience research. However, conventional approaches have several limitations, such as the need for exogenous staining, limited accessibility to volumetric information, and qualitative analysis. Herein, we present high-resolution label-free volumetric imaging and analysis of mouse brain tissue using three-dimensional quantitative phase imaging. Measurement of the refractive index distribution of tissue enables direct imaging of the cellular and subcellular structures. Quantification of subcellular organelles is performed in the anatomical regions of the somatosensory cortex, corpus callosum, caudoputamen, and thalamus regions.

## 1. Introduction

High-resolution structural imaging of brain tissue provides important information for research on neuroscience and the pathophysiology of neurodegenerative diseases (1-3). Recent advances in fluorescence microscopy have enabled sub-diffraction molecular-specific imaging of subcellular organelles and protein distribution (4, 5). Moreover, the combination of tissue-clearing techniques has been extended to volumetric fluorescence imaging (6, 7). However, conventional fluorescence imaging approaches do not directly provide structural information; thus, high-resolution structural information is largely acquired using electron microscopy (EM) (8). Imaging tissue with EM has several limitations, despite its ultra-high spatial resolution imaging ability: (i) the preparation of tissue samples entails a complicated and time-consuming process; (ii) volumetric imaging requires serial sectioning of tissue into thin slices, which usually damages the sample and requires significant time to perform measurements; and (iii) the imaging contrast of EM is not directly related to the electron scattering cross-section in the general EM configuration, and thus does not provide quantitative imaging information.

Herein, we present high-resolution volumetric structural imaging of mouse brain tissue using quantitative phase imaging (QPI). QPI, which exploits the distribution of refractive index (RI) as an intrinsic and quantitative imaging contrast, has enabled label-free imaging and analysis of cells (9). QPI has been employed for research in hematology (10, 11), cell biology (12, 13), immunology, microbiology (14), and regenerative medicine (15, 16). In particular, 2-dimensional (2D) QPI techniques have been utilized for the quantitative analysis of thin tissue sections (17-20). Recently, optical diffraction tomography (ODT), a 3-dimensional (3D) QPI technique based on laser interferometric microscopy and illumination angle scanning has been used for volumetric histopathological analysis (21). However, the use of a coherent light source such as laser inevitably involve speckle noise due to a long coherent length of laser and makes the system vulnerable to environmental vibration or perturbation, which prevents from translating into clinical laboratory. More recently, 3D QPI techniques based on low coherent light sources have been developed (22-25). The use of an incoherent light source significantly reduced the coherent speckle noise while maintain the benefits of QPI.

In this study, we employed a high-fidelity 3D QPI technique that uses a temporally incoherent light source for volumetric imaging and quantitative analysis of mouse brain tissue samples. Systematic 3D imaging was performed for the label-free visualization of thick tissue samples, and also utilized for the quantitative analysis of the subcellular organelles in thick tissues. In particular, the structures of the cerebral cortex somatosensory area layer 1 (CTX-1) and layer 2/3 (CTX-2/3), corpus callosum/caudoputamen (CC/CP), and thalamus (TH) regions are systematically visualized and analyzed. Furthermore, from the measured RI tomograms of thick tissues, their biophysical parameters, including the volumes of the nucleus and nucleoli, dry-mass concentration, and content, were quantitatively analyzed. We envision that the present method will extend the volumetric analysis of tissue samples in various applications, including neuroscience and histopathology.

## 2. Results and Discussion

### 2.1. Volumetric RI measurement of mouse brain tissue using PEPSI-ODT

The RI of mouse brain tissue was measured using plural efficient patterns for self-interfering ODT (PEPSI-ODT) with minimal sample preparation (**Figure 1A**). ODT does not require an external contrast agent for RI measurement because it utilizes the intrinsic optical quantity, i.e., the RI of the sample itself, to provide measurement contrast. This quantitative measurement technique greatly simplifies the sample preparation protocol by eliminating the need for staining to image the structures. PEPSI-ODT reconstructs the 3D RI distribution of a sample using patterned LED illuminations, which enable high-fidelity low-noise imaging.

**Fig. 1.**
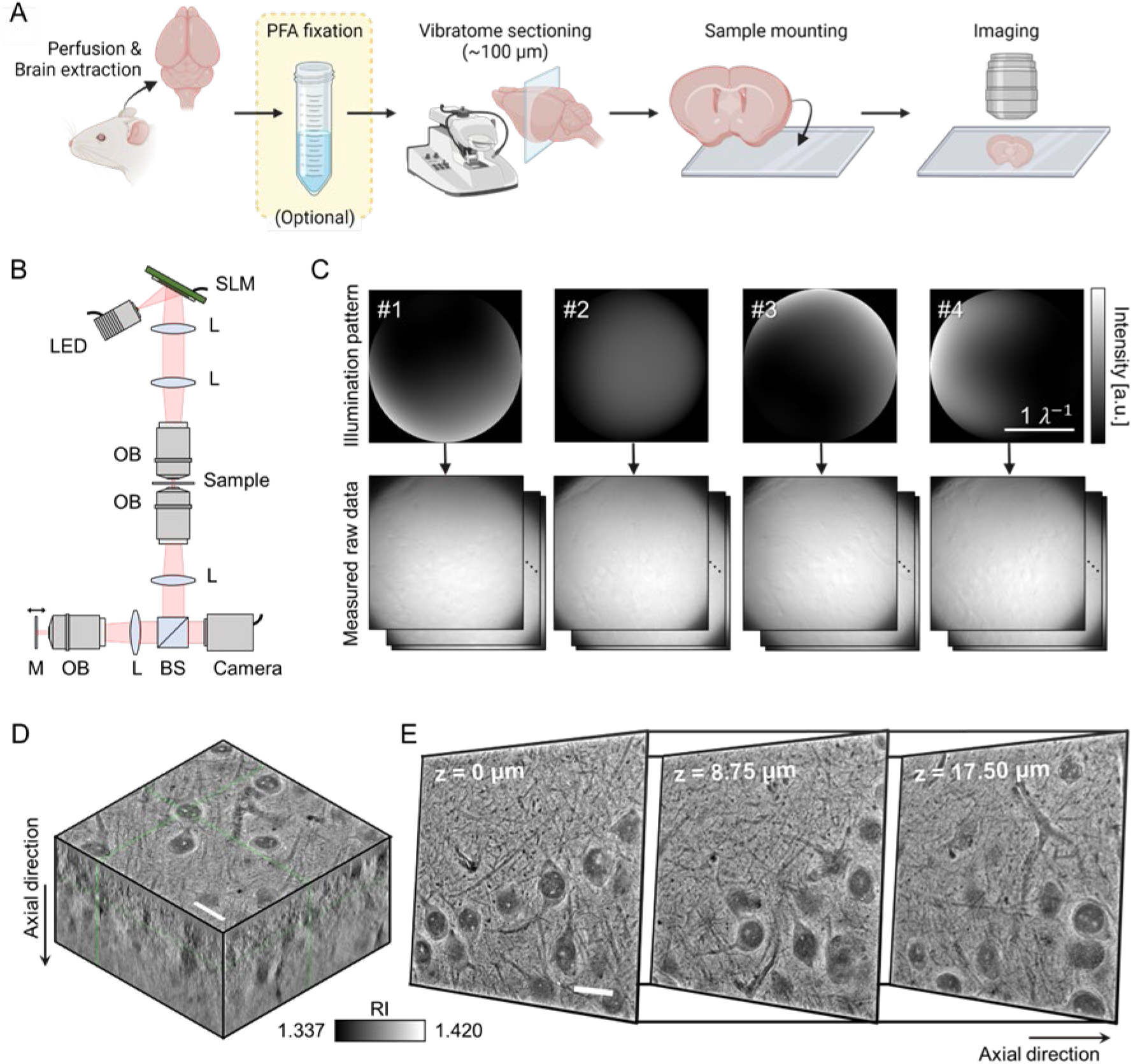
Three-dimensional (3D) refractive index (RI) measurement of mouse brain tissues. **(A) Mouse brain tissue preparation protocol for measurements.** The brain was excised surgically from the mouse after transcardial perfusion of paraformaldehyde (PFA) solution and fixed by immersion in PFA solution to preserve its structure. Immersion fixation is optional but recommended if it is not possible to start the optical measurement immediately after harvesting the brain. The whole brain tissue was then sliced into coronal sections using a vibratome and mounted between two coverslips for imaging. The tissue was mounted with phosphate-buffered saline to maintain physiological conditions. The images were created with BioRender.com **(B) Optical setup for measuring the 3D RI of the sample**. It modulated the illumination intensity patterns in the pupillary plane. The position of the mirror was altered for optical refocusing. **(C) Four illumination intensity patterns are displayed on the SLM**. The patterns were optimized for accurate RI measurement. The corresponding sample image patterns measured in the image plane are shown below. **(D) Reconstructed RI tomogram using the principle of PEPSI ODT**. Scale bar: 20 μm **(E) Each image shown on the sides is the sectional image of the tomogram at three different axes**. The arrow showing the axial direction indicates the direction of the sample depth. Scale bar: 20 μm LED: light emitting diode, SLM: spatial light modulator, L: lens, OB: objective lens, BS: beam splitter, M: mirror, RI: refractive index, ODT: optical diffraction tomography.

Dissected from a mouse, thick brain tissue samples were prepared for measurement by mounting the tissue sections between two glass coverslips. Tissue sections were obtained using a vibratome and sectioned at a thickness of 100 μm. Ideally, it is possible to directly measure the RI tomogram of a thick tissue using this harvesting, sectioning, and mounting procedure. However, these processes often require a specific amount of time which may cause the cells inside the tissue to start deforming and dying. In our experiments, the brain was fixed by perfusion and immersion in paraformaldehyde (PFA) solution (see Materials and Methods), to preserve the cellular organizations of the freshly harvested tissue. However, this fixation protocol is optional and can be skipped for short measurements.

PEPSI-ODT was used to measure the RI tomograms of the brain tissues (**Figure 1B**; see Materials and Methods). The theoretical lateral and axial resolutions of the system were 110 and 350 nm, respectively. A light-emitting diode (LED) was used as a temporally incoherent and spatially low-coherent light source. A spatial light modulator (SLM) was placed in the pupillary plane to modulate the illumination intensity during measurements. The patterns were optimized to maximize spatial frequency coverage and signal strength. The four patterns displayed on the SLM and measured raw intensity image stacks are shown in **Figure 1C**. The measurement focus was altered using a remote focusing scheme. Raw data were utilized to reconstruct the 3D RI tomogram, which used the Rytov approximation under the assumption of a sample with a slowly varying phase distribution (26). A more detailed description of the reconstruction steps is described elsewhere (22).

The reconstructed RI tomogram is shown in **Figure 1D**. Volumetric measurement of the RI was performed using a single-thickness tissue section. 2D sectional visualization of the measurement results at various axial positions revealed the differential organization of various structures inside the tissue without labeling (**Figure 1D**−**1E**). Different cells at various spatial locations were visualized based on the RI values. Subcellular components such as individual cells, nuclear membranes, and nucleoli are clearly visible at sub-micrometer measurement resolution. The blood vessels were also displayed in the RI tomogram, and it is possible to track their positions at various depths.

### 2.2. RI tomograms of mouse brain tissue for various anatomical regions

The RIs of four representative anatomical regions of the mouse brain tissue were measured and analyzed (**Figure 2**) to systematically perform quantitative imaging of mouse brain tissue. Measurements were performed for CTX-1, CTX-2/3, CC/CP, and TH (Figure 2A). In all four regions, the cytoplasmic regions exhibited a lower RI (darker in the image) than that of the outer volume of the cells (Figure 2B). In particular, the nuclear regions of cells had lower RI values than those of the cytoplasm, and the volume within the blood vessels had the lowest RI range (darkest in the image). The nucleoli exhibited the highest RI values. This was highly concurrent with previous studies that reported that cell nuclei have a lower RI and dry mass density than those of the cytoplasm (27). These parameters were studied in various cell cultures, but our study also found a similar tendency in tissues (27).

**Fig. 2.**
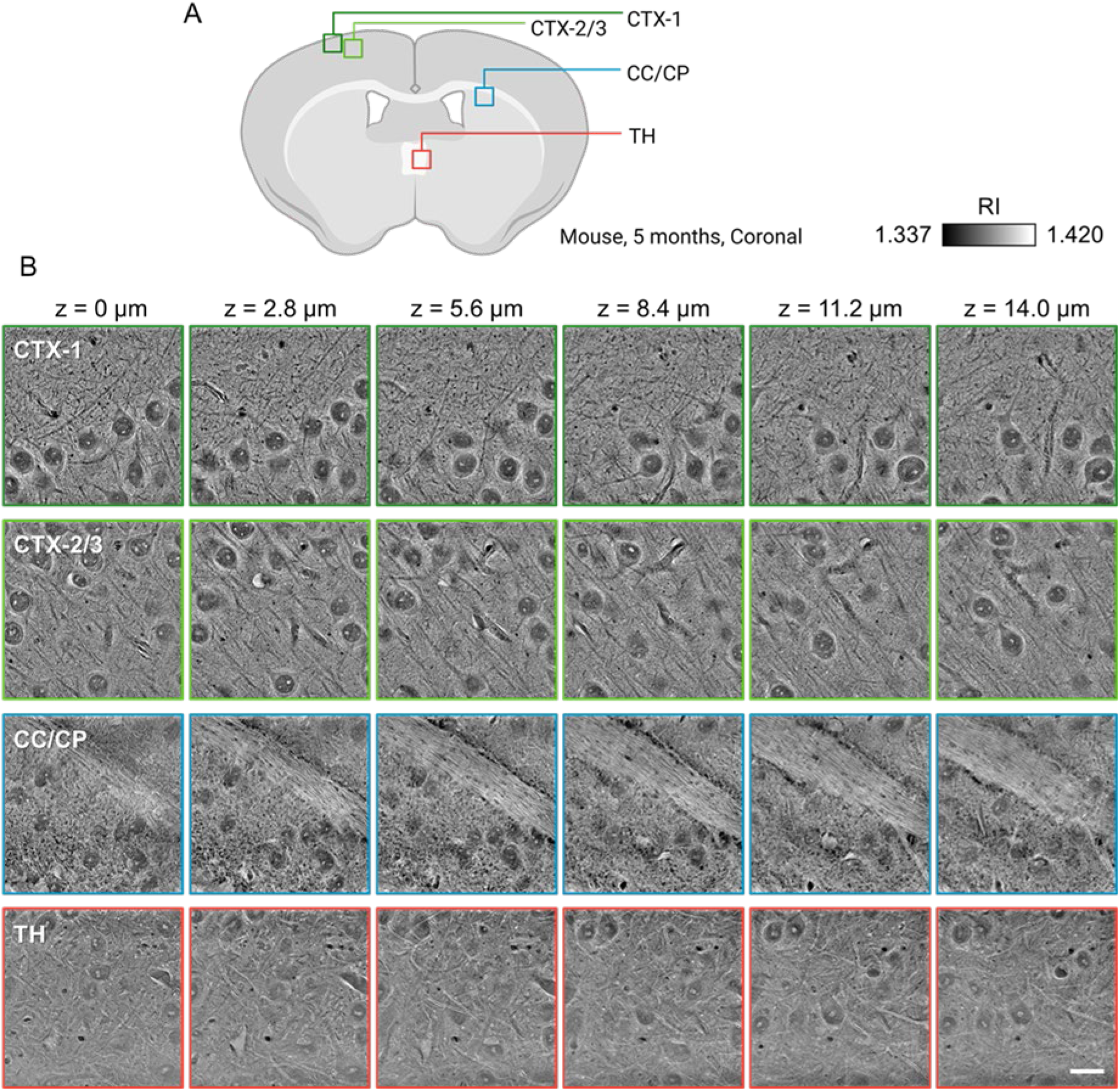
Fig 2. Volumetric measurements of mouse brain tissue at various anatomical regions and depths. **(A) Schematic showing the four different regions of RI measurement in the mouse brain.** The images were created with BioRender.com **(B) Lateral section images of the measured RI tomogram at various depths**. Each region shows different morphology, organization and RI values of the cells and blood vessels. The RI of the nucleus and cytoplasm of the cells was lower than that of the extracellular matrix. Quantitative imaging of the mouse brain tissue with minimal preparation allows volumetric visualization of various structures within the sample. Scale bar: 20 μm. CTX-1: cerebral cortex somatosensory area layer 1, CTX-2/3: cerebral cortex somatosensory area layer 2/3, CC/CP: corpus callosum, TH: thalamus, RI: refractive index.

The RI tomogram shows variations in the positioning and organization of the nuclei in the different layers in the CTX. The CTX-1 region of a coronal section showed multiple nuclei aligned along a line; the nuclei were observed only in the inner area. This arrangement pattern in the CTX-1 area, with two distinct regions with and without nuclei, was maintained throughout the depth of tissue. The deeper region, i.e., CTX-2/3, showed evenly distributed nuclei, whereas the nerve fibers with low RI values showed directionality, which is perpendicular to the brain surface. In comparison, the degree of directionality and alignment of the nuclei and fibrous structures were lesser in the TH. The RI values of the fibers were also different from those observed in the CTX region. The fibers observed predominantly in the TH region were surrounded by a high-RI component with a smaller diameter. A similar structural arrangement was also observed in the CTX region.

A large bundle of nerve fibers was visualized in the CC area with high RI values compared with other areas. Previously, the overall RI of brain tissue slices in the CC region was calculated to be 1.407 ± 0.015 using optical coherence tomography (28). The range of our RI measurement results for the CC, as shown in Figure 2B, was in accordance to the previously reported value. The CC is a white matter region, whose high RI value may be attributed to myelinated axons in the fiber bundle. Myelin sheaths have high lipid content, and a previous study modeled myelin sheaths as alternating spaces with high RI (1.47) and lower RI (1.35) (29). PEPSI-ODT lacks sufficient resolution to depict this alternating structure since it is only a few nanometers thick. However, it is noteworthy that the average RI range of this model is in agreement with our measurements. The RI tomograms obtained by PEPSI-ODT provided quantitative information about the tissue and visualized various tissue components at sub-micrometer resolution.

### 2.3. Volumetric analysis of nuclear morphology in mouse brain tissue

The volumetric images obtained using PEPSI-ODT permit quantitative analysis of the morphology of the nuclei and nucleoli within each nucleus with sub-micrometer spatial resolution (**Figure 3**). Since the RI of biological samples is directly proportional to the protein density, the different internal components can be distinguished based on their RI values. As ODT can measure the RI of samples with sub-micrometer spatial resolution, which is outstanding compared to other RI measurement methods, it can visualize the subcellular components within a given biological sample. ODT is a quantitative imaging technique that offers not only volumetric visualization but also the volumetric RI of a sample.

**Fig 3.**
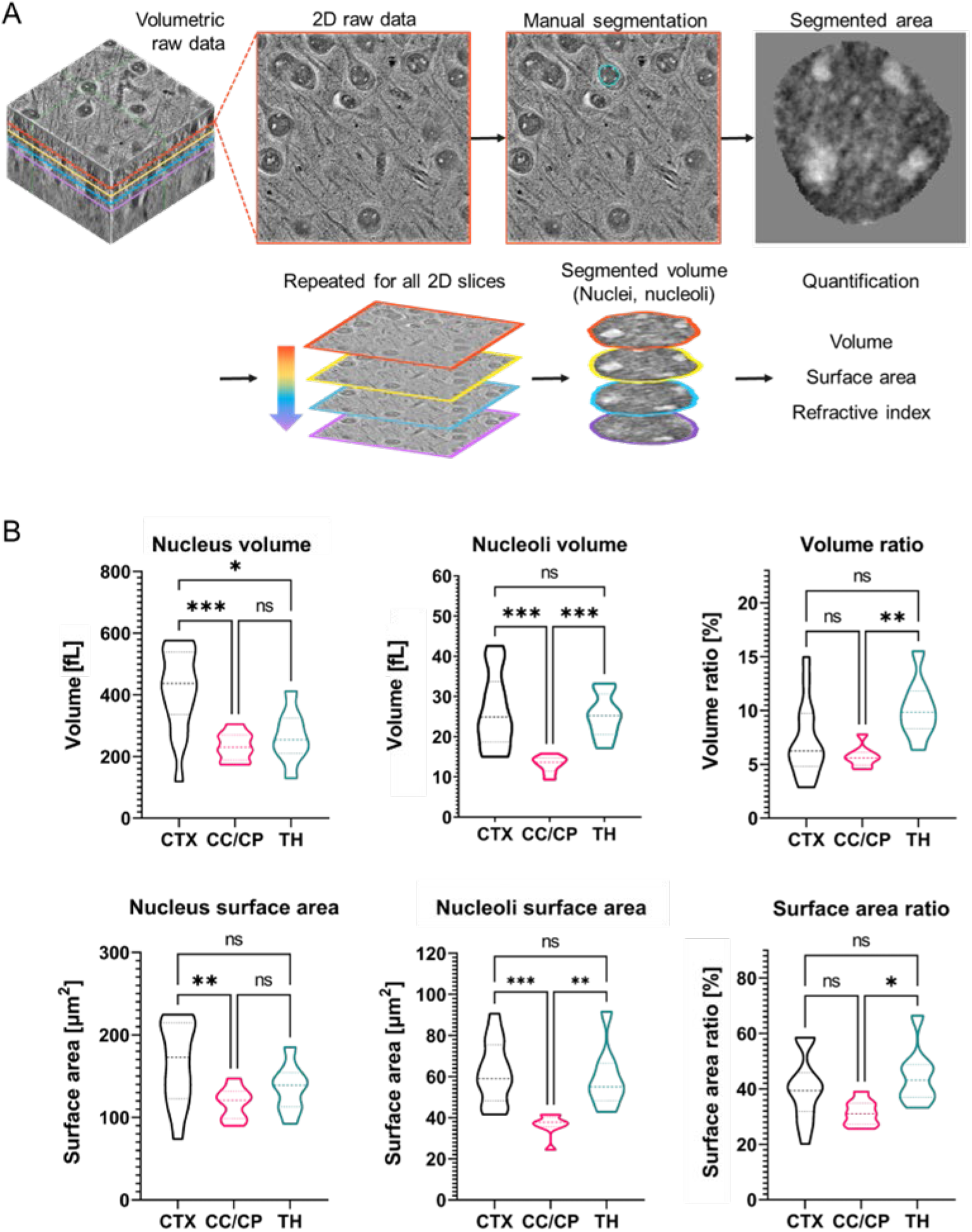
Morphological analysis based on the 3D RI measurements of mouse brain tissue. **(A) Work flow for segmenting individual nuclei and nucleoli inside the tissue**. The raw volumetric data are segmented manually slice-by-slice. The segmented voxels are utilized for further quantitative analysis, such as volume and surface area calculation. **(B) The results of morphological analysis of individual nuclei and nucleoli inside each nucleus are compared among the three different regions in the mouse brain**. The volume and surface area ratios represent the relationship of all nucleoli to the nucleus. CTX: cerebral cortex somatosensory area, CC/CP: corpus callosum, TH: thalamus. The horizontal lines in the graph represent median and quartiles: n = 15 for CTX, n = 8 for CC/CP, and n = 8 for TH.

First, the volumes of the nuclei and nucleoli were segmented from the raw RI tomogram for morphology analysis and dry mass calculation (Figure 3A; see Materials and Methods). Segmentation was conducted manually using MATLAB. The nuclei were segmented based on the clearly visualized boundaries with high RI values (nuclear membrane) for each nucleus, and the high-RI clusters located within were segmented as nucleoli. Clusters with high RI values inside the nucleus have been identified as nucleoli in several previous studies (30). Segmentation was repeated for each 2D slice along the axial direction. The resulting stack of segmented slices was then considered as a single volumetric object, and the volume and surface area of the individual object were calculated. The calculation results for the three different brain regions (CTX, CC/CP, and TH) are shown in Figure 3B. The volume and surface area of the nucleoli represent the values of all nucleoli present in the nucleus. The volume ratio, i.e., the volume of the nucleoli volume divided by the volume of the nucleus, represents the percentage volume occupied by all the nucleoli present inside a single nucleus. The surface areas were analyzed in a similar manner. A comparison of the volume and surface area in different brain areas revealed that the volume and surface area of the cells in the CTX were greater than those in other areas. In contrast, the volume and surface area of the nucleoli in the cells in the CC/CP were significantly smaller. The volume and surface ratio values were the largest for cells in the TH.

### 2.4. RI distribution and dry mass analysis in various brain regions

Histograms of the RI values within the segmented voxels of the nucleus and nucleoli in the RI tomogram were compared in several brain regions (**Figure 4A**). The minimum RI values of the nuclei were almost similar in all three regions, which approximated 1.350. However, the maximum RI values of the nuclei were considerably lower in the CTX region than those in the CC and TH regions. In all three regions, the RI histogram of the nucleoli was located at the higher tip of the RI histogram of the nuclei.

**Fig 4.**
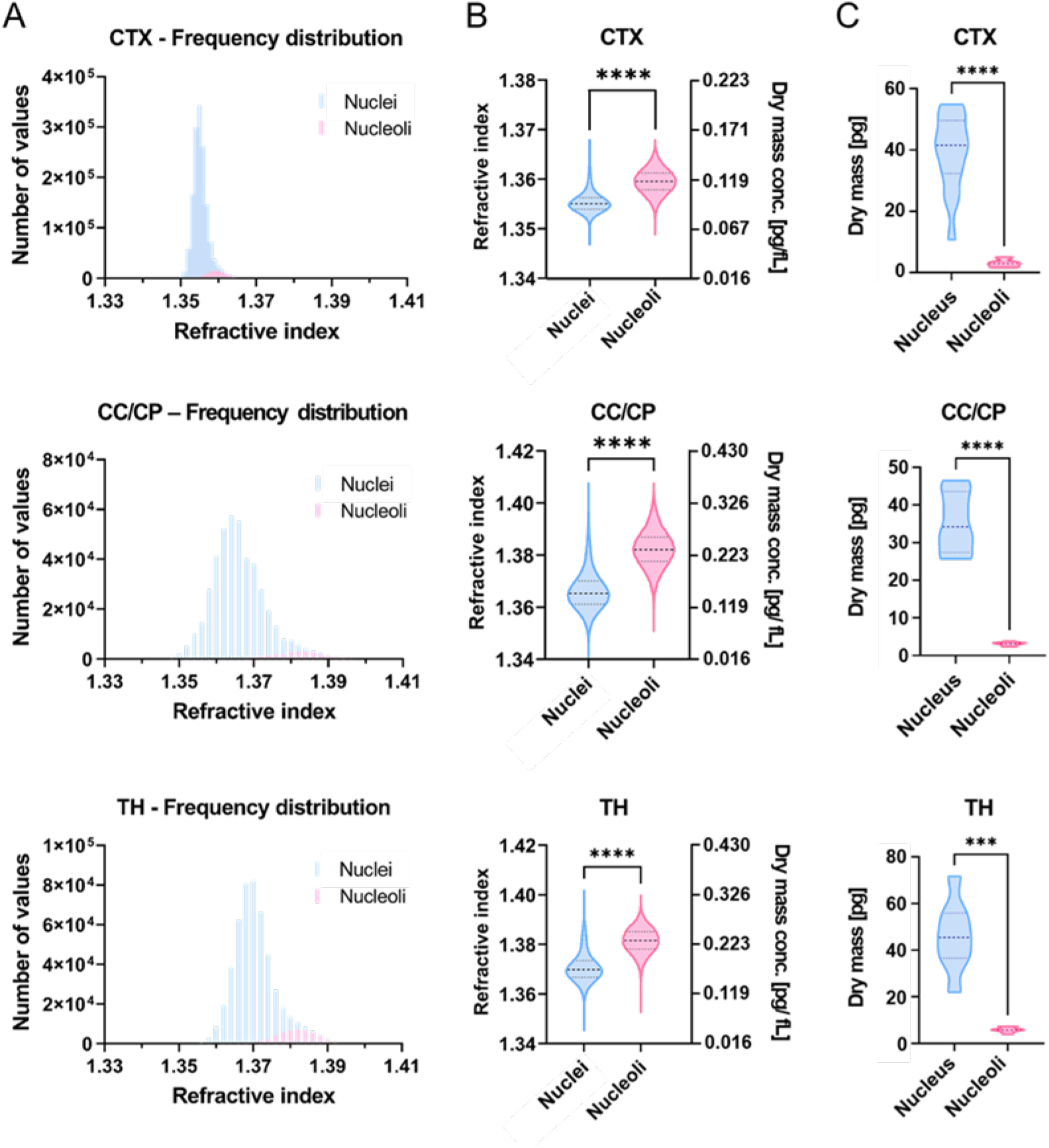
Characterization of RI tomograms measured at various anatomical regions of the mouse brain. **(A) Frequency distribution of the RI values of the nuclei and nucleoli in different regions inside the mouse brain tissue.** The bin size of each histogram is 0.001 for the CTX and 0.002 for the CC/CP and TH each. **(B) RI and dry mass concentration distribution of the nuclei and nucleoli in mouse brain tissue**. The RI values are converted into dry mass concentration using a specific refractive increment of 0.193 fL pg^-1^. The horizontal lines in the graph represent median and quartiles: N_nuclei_=1465248 and N_nucleoli_=93366 for CTX, N_nuclei_=441645 and N_nucleoli_=24841 for CC/CP, and N_nuclei_=498324 and N_nucleoli_=47605 for TH. **(C) Dry mass distribution of the nuclei and nucleoli inside each nucleus in mouse brain tissue**. The horizontal lines in the graph represent median and quartiles: n=15 for CTX, n=8 for CC/CP, and n=8 for TH. CTX: cerebral cortex somatosensory area, CC/CP: corpus callosum, TH: thalamus.

The RI is an important and useful physical parameter because it reflects the local dry mass concentration. This ability of quantitative measurement, which confers a significant advantage while imaging tissues using PEPSI-ODT, originates from the non-specificity of light refraction. The relationship between the RI values and dry mass concentration is represented as follows (31):

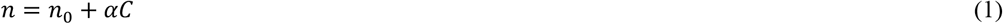

where *n* is the RI of the target substance, *n*_0_ is the RI of the medium, and *C* is the dry mass concentration of the target substance. Here, *α* denotes the specific refractive increment, which is a constant equaling 0.0193 fL pg^-1^. Previous studies have found this value to be practically independent of the composition of various molecules that commonly exist in the cells (32, 33).

In our study, 0.0193 fL pg^-1^ was adopted as the specific refractive increment, and the RI of the medium was set to 1.337. Using this relationship, the dry mass concentration was calculated for each voxel inside the tomograms. The measurement results for the different brain tissue regions are shown in **Figure 4B**. The combination of the dry mass concentration with the spatial information of each nucleus and nucleolus yielded the dry mass of each object (**Figure 4C**). The dry mass of each nucleus and nucleolus were characterized and compared among different brain regions. In all three regions, the dry mass concentrations were significantly higher in the nucleoli, but the total dry mass was greater in the nuclei due to their larger volumes.

## 3. Conclusion

Herein, we described high-resolution volumetric structural imaging of mouse brain tissue and quantitative analysis of subcellular organelles. A 3D RI tomogram of the tissue samples was reconstructed with a sub-micrometer spatial resolution using PEPSI-ODT. The present approach allows label-free imaging as well as analytical methods for the study of brain tissues, which encompass multiscale spatial resolution, ranging from hundreds of nanometers to hundreds of micrometers in volumetric spaces. RI tomograms of the subcellular tissue structures in the anatomical regions CTX-1, CTX-2/3, CC/CP, and TH of the brain were measured, and their biophysical parameters, including the volumes of the nucleus and nucleoli, dry mass concentration, and content, were analyzed quantitatively. We emphasize that the present approach can provide quantitative analysis of subcellular organelles, including the volumes, surface areas, dry mass content, and dry mass concentrations of the nuclear membranes and nucleoli in each anatomical region. This is because the RI value is linearly proportional to the concentration of biomolecules. Minimal tissue preparation for imaging is another substantial benefit of label-free measurements using PEPSI-ODT.

The retrieved and analyzed subcellular organelle information of brain tissue can be studied while researching the pathophysiology of neurological diseases. For example, enlarged and prominent nucleoli are well-known histopathological indicators of cancer. The morphology of the nucleolus is strongly correlated with various neurodegenerative diseases such as amyotrophic lateral sclerosis and frontotemporal dementia. We presented in-depth volumetric and quantitative analyses of the nuclei and nucleoli, which can also be further investigated in the context of liquid-liquid phase separation (30, 34).

Although the present method enabled precise and quantitative morphological analysis of brain tissue, RI distribution does not directly provide molecule-specific information. For correlation analysis of the morphology- and molecular-specific information, one may consider the combination of ODT with fluorescence imaging (35-38). Recently developed machine learning-enabled computational molecular specificity can also be applied to RI tomography of brain tissues (13, 39). Furthermore, the present approach can provide complementary and synergistic information in addition to various spatial biology approaches that exploit DNA, RNA, or protein expression levels at single-cell resolution. Machine-learning-enabled cell type classification methods can also be applied to the reconstructed 3D RI tomogram of brain tissues so that various cell types in the brain can be identified without labeling. We envision that the present approach can unlock new research and diagnostic opportunities in neurology and neurosurgery.

## 4. Experimental Section

### Preparation of Mouse Brain Tissues

Animal care and experimental procedures were performed under approval from the Institutional Animal Care and Use Committee (IACUC) of the Korea Advanced Institute of Science and Technology. All experimental procedures in this study were performed in accordance with the IACUC-approved guidelines. C57BL/6 mice (male, 5 months old, housed with unrestricted access to food and water) were anesthetized by intraperitoneal injection of avertin (2,2,2-tribromoethanol; T48402, Sigma Aldrich) at a volume of 20 μL g^−1^ based on body weight of the mice. Transcardial perfusion was performed in live animals. 20 mL of ice-cold phosphate-buffered saline (PBS) and 20 mL of ice-cold 4% PFA in PBS were perfused consecutively at a rate of 5 mL min^−1^ to wash out the blood and fix the tissue. The brain tissue was carefully excised surgically after perfusion. For complete fixation of the brain tissue, the sample was immersed in 35 mL of PFA solution with gentle shaking at 4 °C at room temperature for 12 h. The sample was then washed with 35 mL of PBS by gently shaking it at 4 °C and room temperature for 12 h each. In ice-cold PBS to minimize possible damage, the brain tissue was sectioned into 100-μm-thick slices using a vibratome (VT 1200S, Leica) at a speed of 0.3 mm s^−1^ and an amplitude of 1.2 mm s^−1^. The brain tissue section was placed between two #1 glass coverslips (length × width: 24 × 50 mm; Matsunami) using spacers. PBS was used as the mounting medium.

### 3D RI Measurement Optical System

A custom-built optical imaging system was utilized for measuring the 3D RI tomogram (Fig. 1B). This ODT system utilizes an LED (M470L2, Thorlabs) as an incoherent light source and an SLM (X10468-01, Hamamatsu) to generate illumination intensity patterns. In contrast to conventional ODT systems that utilize a coherent plane wave to illuminate the sample at various angles, we retrieved the RI value of the sample using Köhler illumination and placing the spatial light modulator in the pupillary plane to modulate the intensity. The four illumination intensity patterns were optimized to achieve accurate RI value retrieval, also known as PEPSI-ODT. The intensity-modulated light was then transmitted through the sample and two objective lenses (LUMFLN60XW, NA=1.1, Olympus). The light was then split using a beam splitter and reflected by a mirror on a piezo stage (LPS710E/M, Thorlabs) to refocus the beam using a remote focusing scheme. The resulting raw data were recorded with a camera (LT425M-WOCG, Lumenera Inc.) and deconvolved using the point spread function of the illumination pattern to obtain RI. More detailed information on PEPSI-ODT and its differences from other types of ODT can be found elsewhere.(22) The RI values in the tomogram obtained using PEPSI-ODT were calibrated to have the minimum RI value of the medium, i.e., 1.337.

### Quantification of Nuclei and Nucleoli

The volumes of the nuclei and nucleoli were manually segmented from the RI tomogram using MATLAB. The volume of each nucleus or nucleolus was obtained by adding the areas segmented at each 2D x-y plane section in the 3D tomogram. After defining the volume of each object, the morphological properties, such as volume and surface area, were calculated. The physical dimensions of each voxel were 0.11 μm in the lateral directions and 0.35 μm in the axial direction. The nucleus included the nucleoli, and the volume and surface area of the nucleoli described in Figure 3 are the sum of all the nucleoli present inside a single nucleus. Thus, the volume and surface area ratio indicate the volume or surface area of all the nucleoli relative to the nucleus. The RI values of the voxels inside each object were used to calculate the frequency distribution and dry mass concentration. The dry mass concentration C is calculated using Equation (1), where the medium RI value, n_0_ was 1.337, and the specific refractive increment α was 0.193 fL pg^−1^. The dry mass of each nucleus and the nucleoli inside the nucleus were calculated by summation of the dry mass values obtained at each voxel. The dry mass of each voxel was calculated by multiplying the dry mass concentration by the voxel volume.

### Statistical Analysis

Frequency distribution analysis and statistical tests were conducted using GraphPad Prism version 9.1.2 for Windows (GraphPad Software, San Diego, California USA, www.graphpad.com). Statistical significance was set at p < 0.05.

The distribution of the volume and surface area data in Figures 3 and 5 show normal distribution (passed the Anderson-Darling test, D’Agostino & Pearson test, Shapiro-Wilk test, and Kolmogorov-Smirnov test, alpha=0.05), except for the nucleoli surface area data in the CC/CP in Figure 3. As shown in Figure 3, multiple comparisons between different anatomical regions were performed using the Kruskal-Wallis test for the nucleoli surface area data and Brown-Forsythe and Welch ANOVA tests for the other data. As shown in Figure 5, comparisons between the groups were performed using an unpaired t-test with Welch’s correction.

The RI data in Figures 4 and 5 exhibited non-normal distribution (did not pass the Anderson-Darling test, D’Agostino and Pearson test, and Kolmogorov-Smirnov test, alpha=0.05). Comparisons between the groups were performed using the Mann-Whitney U test with Hodges-Lehmann difference calculation. The dry mass data in Figures 4 and 5 showed normal distribution (passed the Anderson-Darling test, D’Agostino and Pearson test, Shapiro-Wilk test, and Kolmogorov-Smirnov test, alpha=0.05). Comparisons between the groups were performed using an unpaired t-test with Welch’s correction.

## Acknowledgements

This work was supported by National Research Foundation of Korea (2015R1A3A2066550, 2021R1A2C2005294, 2022M3H4A1A02074314), Institute of Information & communications Technology Planning & Evaluation (IITP; 2021-0-00745) grant funded by the Korea government (MSIT), KAIST Institute of Technology Value Creation, Industry Liaison Center (G-CORE Project) grant funded by MSIT (N11230131), and the Korea Health Technology R&D Project through the Korea Health Industry Development Institute (KHIDI), funded by the Ministry of Health & Welfare, Korea (HI21C0977).

